# Cancer-cell derived S100A11 promotes macrophage recruitment in ER+ breast cancer

**DOI:** 10.1101/2024.03.21.586041

**Authors:** Sanghoon Lee, Youngbin Cho, Yiting Li, Ruxuan Li, Daniel Brown, Priscilla McAuliffe, Adrian V Lee, Steffi Oesterreich, Ioannis K. Zervantonakis, Hatice Ulku Osmanbeyoglu

## Abstract

Macrophages are pivotal in driving breast tumor development, progression, and resistance to treatment, particularly in estrogen receptor-positive (ER+) tumors, where they infiltrate the tumor microenvironment (TME) influenced by cancer cell-secreted factors. By analyzing single-cell RNA-sequencing data from 25 ER+ tumors, we elucidated interactions between cancer cells and macrophages, correlating macrophage density with epithelial cancer cell density. We identified that S100A11, a previously unexplored factor in macrophage-cancer crosstalk, predicts high macrophage density and poor outcomes in ER+ tumors. We found that recombinant S100A11 enhances macrophage infiltration and migration in a dose-dependent manner. Additionally, in 3D models, we showed that S100A11 expression levels in ER+ cancer cells predict macrophage infiltration patterns. Neutralizing S100A11 decreased macrophage recruitment, both in cancer cell lines and in a clinically relevant patient-derived organoid model, underscoring its role as a paracrine regulator of cancer-macrophage interactions in the pro-tumorigenic TME. This study offers novel insights into the interplay between macrophages and cancer cells in ER+ breast tumors, highlighting S100A11 as a potential therapeutic target to modulate the macrophage-rich tumor microenvironment.

## Introduction

The complex breast tumor microenvironment (TME) exhibits heterogenous cell type composition patterns across breast cancer subtypes [1, 2]. Macrophages represent an abundant immune cell type in the TME and have been shown in preclinical studies to promote tumor growth and metastasis [3–10]. Clinically, estrogen receptor-positive (ER+) breast tumors with a high macrophage density exhibit poor response rates to hormone therapy [11, 12]. Hence, there is a strong interest in uncovering cancer-macrophage interactions that predict disease outcomes and develop therapeutic strategies to target the pro-tumorigenic macrophage-rich ER+ breast TME.

Previous studies have shown that chemokines CCL2 and CCL5 promote macrophage infiltration in ER+ breast cancer [13]. The expression levels of these chemokines are regulated by estrogen [13]. Treatment of human breast tissue explants with tamoxifen reduced macrophage trafficking, whereas *in vivo* xenografts exposed to estradiol exhibited increased macrophage density [13]. Another study showed that CCL2 is regulated by the activation of NF-κB signaling in ER+ breast cancer cells, demonstrating that proinflammatory signaling pathways represent candidate targets to limit macrophage infiltration [14]. However, these studies did not investigate cancer-macrophage crosstalk in a physiologically relevant 3D microenvironment and only focused on the role of the CC chemokine ligand family (CCL) in macrophage recruitment.

Single-cell RNA-seq (scRNA-seq) is a powerful tool for investigating cellular heterogeneity, paracrine cell-cell interactions, and pathways enriched at the single-cell level in the complex breast tumor TME. A landmark study on treatment-naïve breast cancers demonstrated how single-cell transcriptional features can be utilized to predict clinical outcomes by identifying distinct tumor ecosystem classes [15]. Another study utilized scRNA-seq analysis of pre-and post-treatment biopsies to demonstrate how macrophage interactions with T cells lead to differential responses to immune-checkpoint therapy [16]. Furthermore, spatial omics approaches provide complementary information on cell-cell crosstalk in the native tumor microenvironment [17–20]. For example, Onkar et al. [21] recently revealed that macrophage-T cell-interacting neighborhoods have prognostic value in treatment-naïve ER+ breast cancer. Notably, among all immune cell types, macrophages exhibit the highest infiltration in the ER+ breast tumor microenvironment [21]. Therefore, new approaches that employ single-cell technologies with functional perturbation studies hold promise for identifying potential therapeutic targets in the macrophage-rich ER+ breast tumor microenvironment.

Here, we investigated cancer cell-derived factors that promote macrophage recruitment in ER+ breast cancer by combining scRNA-seq analysis with experimental validation in a 3D cancer-macrophage co-culture. We found that tumors with high S100A11 expression in cancer cells exhibited a high macrophage density. Furthermore, bulk transcriptomic analysis revealed that ER+ breast cancer tumors overexpressing S100A11 are associated with worse survival outcomes. Using multiple tumor models, including established ER+ breast cancer cell lines and patient-derived organoids, we demonstrated that S100A11 neutralization limits macrophage recruitment in a 3D microenvironment. Consistent with these results, the treatment of human primary macrophages with S100A11 promoted cell motility and 3D infiltration in the absence of cancer cells. Our systems biology approach presents a powerful strategy for the discovery of targetable cancer-macrophage paracrine mechanisms that establish a pro-tumorigenic macrophage-rich microenvironment.

## Results

### A single-cell cell atlas for ER+ breast tumors

We integrated two treatment-naïve ER+ breast cancer single-cell RNA sequencing (scRNA-seq) datasets obtained from Wu et al. (n=10) [15] and Bassez et al. (n=15) [16], resulting in a combined dataset of 72,079 cells from 25 tumors (**Fig. 1A and Fig. 1B**). Details regarding the breast tumor subtype composition and quality control metrics are provided in **Supplementary Table 1**. We conducted principal component analysis (PCA) using the top 2,000 variably expressed genes across all cells. The cells were classified into transcriptionally distinct clusters based on the top 30 principal components (**Fig. 1B; Supplementary Fig. 1A**). Notably, the clustering of scRNA-seq samples, determined by nearest neighbors, did not align with clustering by patient or study, suggesting the successful mitigation of batch effects.

**Figure 1.**
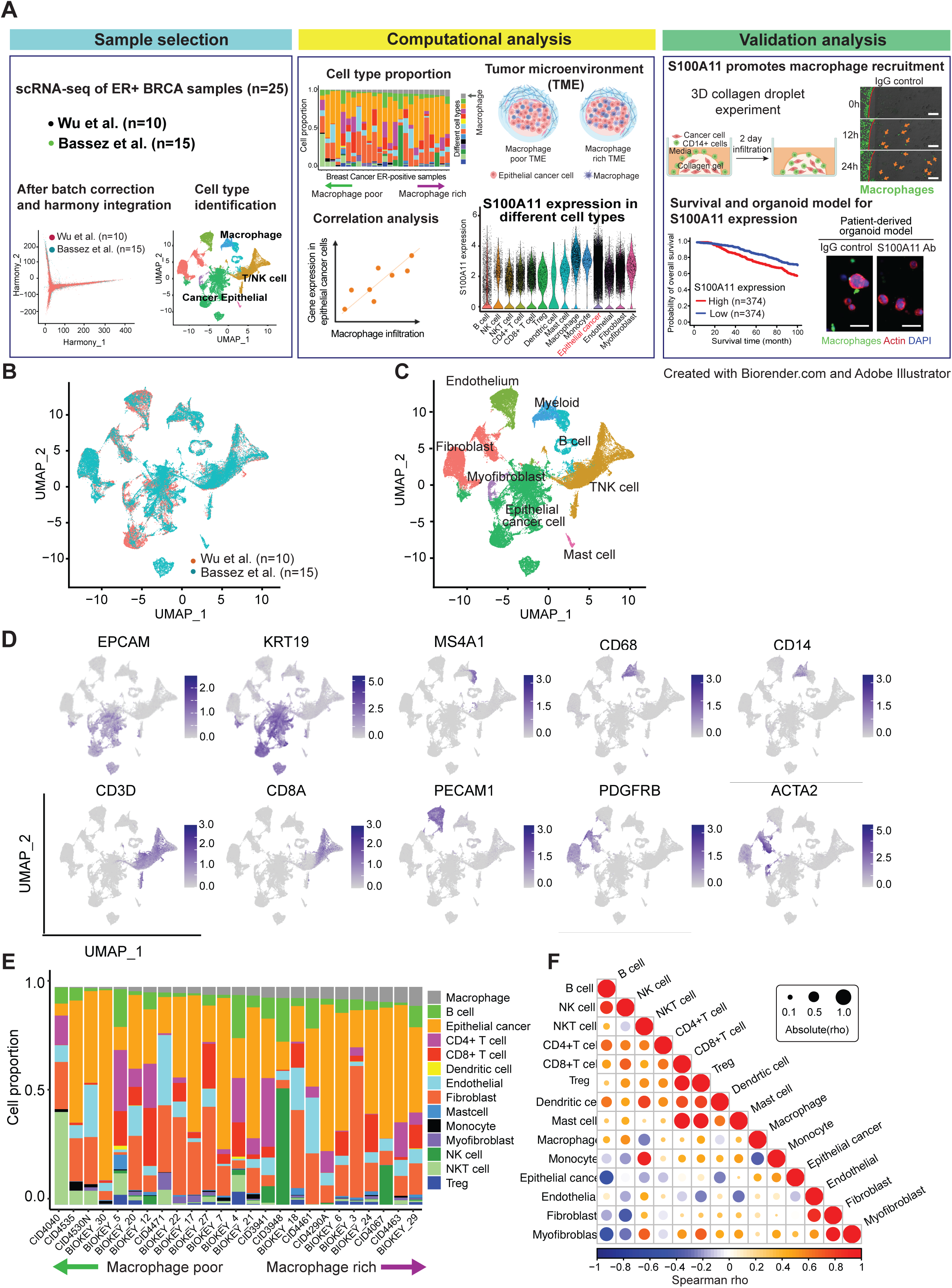
Characterization of ER+ breast cancer tumor microenvironment and cell types. (A) Summary of the data integration and analysis workflow. A total of 72,079 cells analyzed by integrated scRNA-seq across 25 ER+ primary breast tumors. The expression of S100A11 in cancer epithelial cells was validated in BRCA cell lines and organoid models. (B) Uniform manifold approximation and projection (UMAP) clustering of the integrated scRNA-seq data colored by datasets and **(C)** by cell types. **(D)** Feature plots showing the expression levels of selected canonical cell markers that are used to identify the clusters for each cell type. More detailed cell markers are described in Methods. **(E)** Stacked bar plot showing the proportion of endothelial, immune, stromal, and tumor cell types in individual tumors relative to the total cell count. The 25 tumors in x-axis were ordered from macrophage-poor on the left to macrophage-rich on the right. **(F)** Spearman correlation pyramid plot for the 14 cell type proportions. macrophage proportion has positive correlation with ECCs but negative correlation with NKT cells and monocyte proportions.

Utilizing the integrated scRNA-seq dataset and canonical marker genes, we conducted comprehensive quantification of 14 primary cell types (**Fig. 2C and D**). Immune cell types include macrophages, dendritic cells (DCs), T cells (CD4^+^ and CD8^+^ T cells), regulatory T cells (Tregs), B cells, natural killer (NK) cells, NKT cells, monocytes, and mast cells. Among non-immune cells, epithelial cancer cells (ECCs), endothelial cells, fibroblasts, and myofibroblasts were identified (**Fig. 2D; Supplementary Fig. 1B and C**). The clustering of immune, tumor, and stromal cells was based on cell identity rather than on patient origin. Consequently, our ER+ breast cancer atlas incorporates a total of 72,079 single cells annotated into 14 main cell types, including 27,014 cancer cells and 25,154 immune cells.

**Figure 2.**
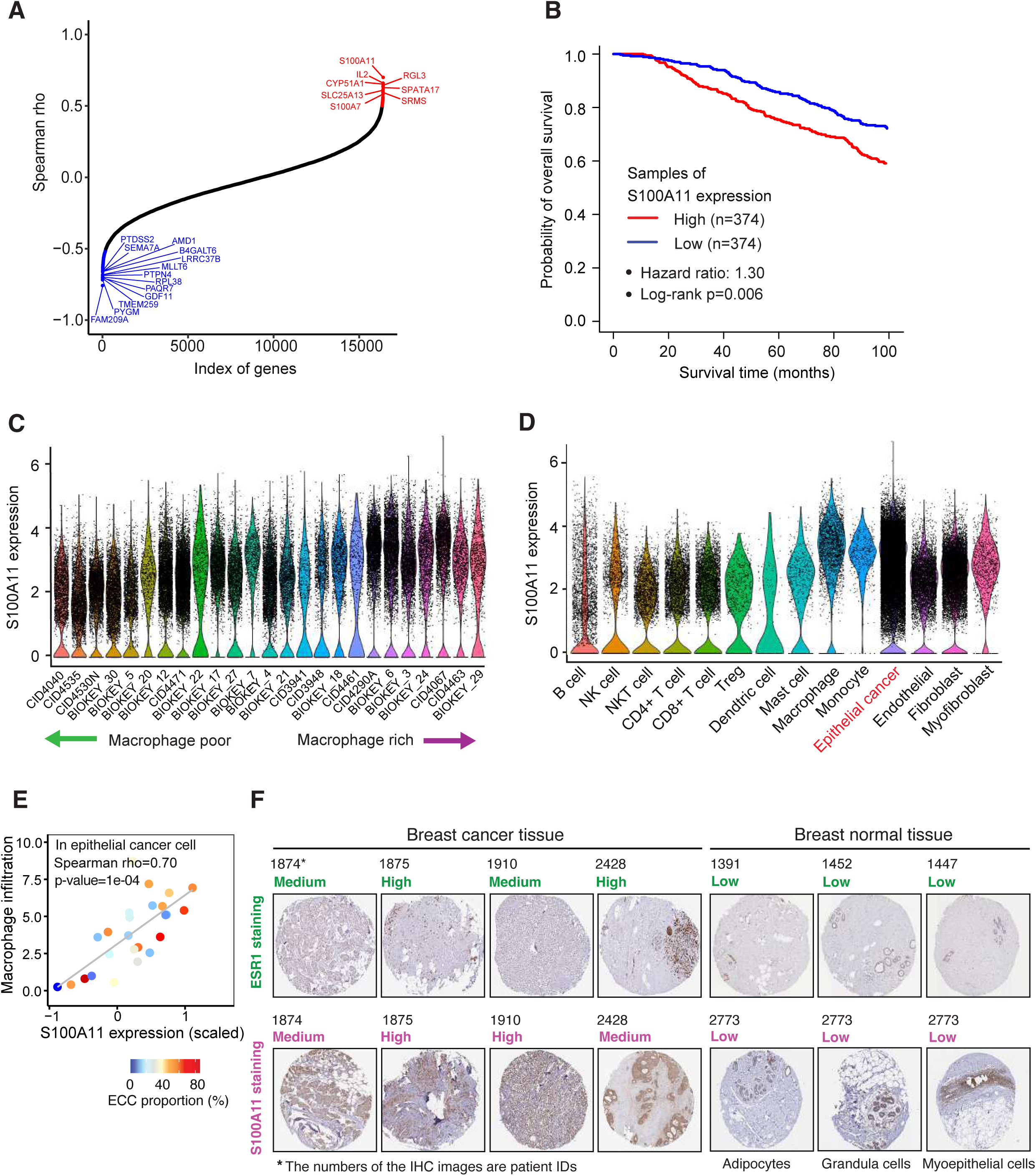
S100A11 expression in single cells and its correlation with macrophage infiltration. **(A)** Line dot plots revealing the correlation between whole gene expression and macrophage infiltration in ECCs or in macrophages. The genes with Spearman correlation abs(rho) > 0.5 were colored in red for positive and blue for negative correlation. The gene symbols of abs(rho) > 0.6 were annotated. **(B)** Survival plot showing the high S100A11 expression in ER+ breast cancer samples is associated with a lower overall survival rate (hazard ratio 1.30 and log-rank p-value 0.006). (C) Violin plot showing the S100A11 expression in individual samples when the samples are sorted by macrophage infiltration in ascending order from left to right. Spearman rho between macrophage infiltration and S100A11 expression in ECCs is 0.7. **(D)** Violin plot displaying the S100A11 expression levels in the different 14 cell types. **(E)** Dot plots showing the correlation between S100A11 expression and macrophage infiltration in ECCs. The mean of the S100A11 expression and macrophage infiltration were calculated in each sample and the color of each dot represents ECC proportion in individual samples. **(F)** Immunohistochemical staining for ESR1 and S100A11 protein expression in breast cancer or normal breast tissue. In breast cancer tissue, the expression of ESR1 and S100A11 proteins were observed in the matched samples.

Next, we determined the macrophage proportion in each tumor and arranged the samples based on the macrophage proportion, ranging from low (macrophage-poor) to high (macrophage-rich) (**Fig. 2E; Supplementary Fig. 1D**). We observed a weak positive correlation between the proportion of macrophages and the proportion of epithelial cancer cells (ECCs) because of the small number of samples (Spearman correlation rho = 0.16) (**Fig. 2F**).

### Transcriptional predictors of macrophage-rich ER+ breast tumors

To identify cancer cell-driven predictors of macrophage infiltration, we computed Spearman rank correlations (rho) between the macrophage proportion and mean gene expression in cancer cells for each tumor sample. We have identified both novel and known relationships. Of the top eight genes whose high expression was associated with a macrophage-rich TME, only three genes have been previously characterized to be expressed in the extracellular domain (secreted factors), including S100A11, RGL3, and S100A7 (**Fig. 2A and Supplementary Table 2**). RGL3 (<u>R</u>al<u>G</u>ef-like-3) is a member of the Ral guanine nucleotide exchange factor (RalGEFs) family that serves as a downstream effector for both Rit and Ras, which can induce oncogenic transformation when they are constitutively active [22]. However, its function in breast cancer and its connection with macrophages remain unexplored. S100A11 and S100A7 belong to the S100 family of calcium-binding proteins and are known to regulate cell migration and proliferation via cytoskeleton remodeling in multiple cancer types [23–28].

To evaluate clinical relevance, we explored the association of these genes with outcomes in ER+ breast tumors using the Molecular Taxonomy of Breast Cancer International Consortium (METABRIC) dataset [29]. The expression of RGL3 and S100A7 was not associated with overall survival outcomes (hazard ratio = 0.95, log-rank test p-value = 0.614 for RGL3; hazard ratio = 1.03, log-rank test p-value = 0.786 for S100A7) (**Supplementary Figure 2A**). In contrast, high S100A11 expression was associated with a poor survival outcome in ER+ breast tumors, with a hazard ratio (HR) of 1.30 (**Fig. 2B**, log-rank test p-value = 0.006). RGL3 and S100A7 showed no predictive value as potential therapeutic vulnerabilities; therefore, we opted not to pursue experimental investigation for RGL3 and S100A7. Instead, we focused on further exploring the association between secreted factor S100A11 and macrophage recruitment.

Analysis of S100A11 expression at the single-cell level confirmed that cancer cells exhibited high expression (**Fig. 2C-E)**. We next compared immunohistochemically (IHC) stained images for S100A11 between ER+ breast cancer tissues and normal breast tissues obtained from the Human Protein Atlas (HPA) database [30]. IHC images demonstrated that S100A11 protein was highly expressed in ER+ breast cancer tissues, but was not detected in normal breast tissues (**Fig. 2F**). Collectively, these survival results and the overexpression in tumors compared to normal tissues suggest that S100A11 represents a potentially actionable target in the ER^+^ breast tumor microenvironment.

### S100A11 enhances macrophage migration speed and recruitment in 3D matrices

We next employed cultures of primary human macrophages to investigate the role of exogenous S100A11 in macrophage migration and infiltration in a 3D microenvironment. Using time-lapse imaging, we evaluated the migration trajectory and speed of primary PBMC (peripheral blood monocyte)-derived macrophages exposed to recombinant S100A11 (1 and 10ng/ml). We found that S100A11 treatment increased migration speed in a dose-dependent manner (**Fig. 3A-B**). To further evaluate the role of S100A11 in macrophage migration in a 3D extracellular matrix, we used an inverted migration assay. The number and position of macrophages inside the 3D environment were tracked following the establishment of a concentration gradient of S100A11 (**Fig. 3C**). Consistent with the migration speed results, the number of recruited macrophages increased in an S100A11 dose-dependent manner compared to that in the control medium (**Fig. 3C**). Taken together, these results demonstrate that soluble S100A11 potentiates both the macrophage migration speed and macrophage infiltration in a 3D extracellular matrix.

**Figure 3.**
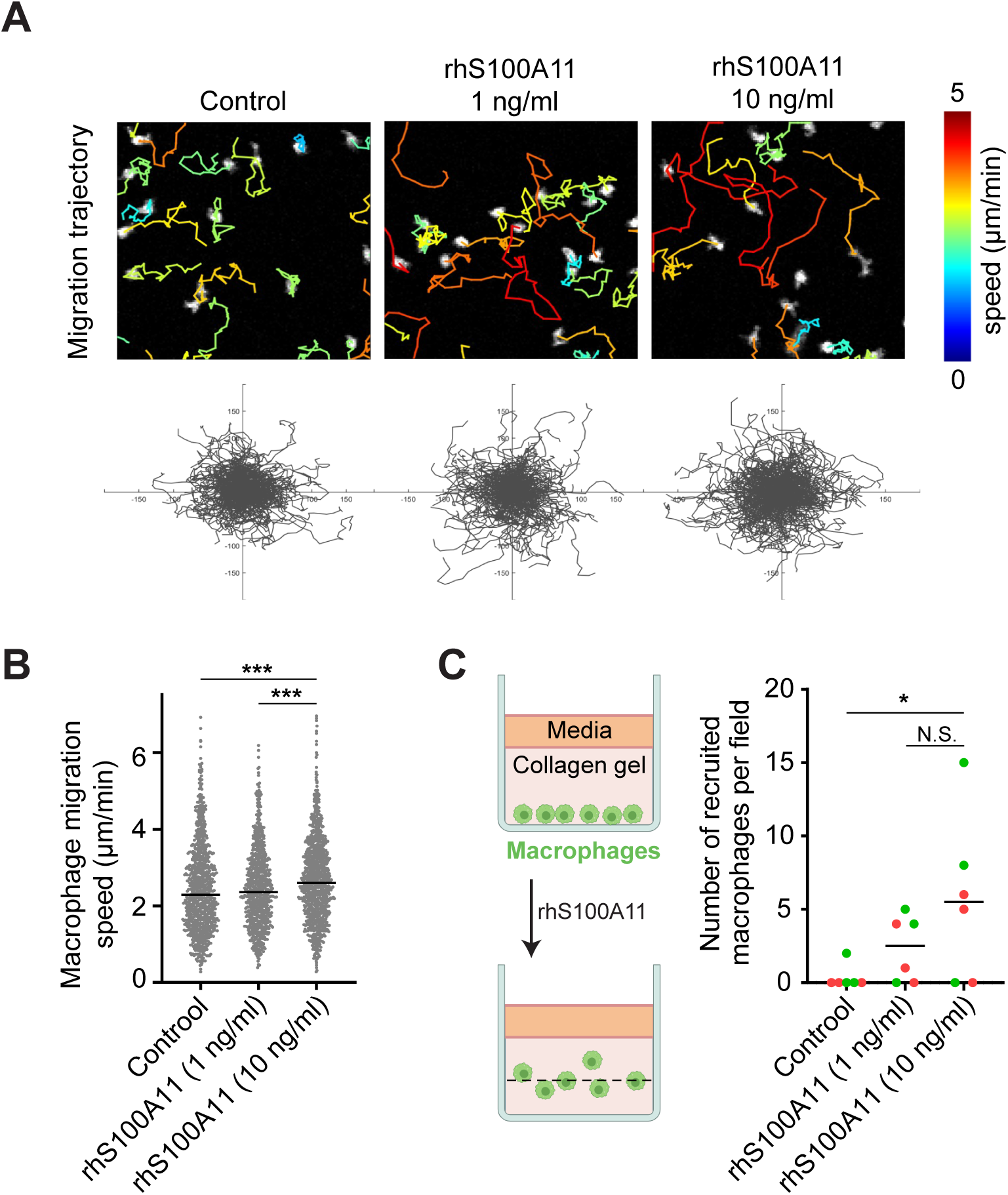
S100A11 protein induces migratory phenotype of macrophages and promotes recruitment in collagen matrix. **(A)** (Upper) Human macrophage migration trajectory and average instantaneous speed is overlaid on the FITC stained macrophages under the treatment of rhS100A11. (Down) Rose plot of macrophage trajectory over 1 h. Macrophage migration was imaged every 2 min. (n>237 tracks for each condition). **(B)** Quantification of average instantaneous migration speed of macrophages under the treatment of rhS100A11. S100A11 increased macrophage migration speed. (n>237 tracks). **(C)** Quantification of macrophage recruitment in collagen matrices using the inverted invasion assay. Number of macrophages recruited to 20 µm above the surface was analyzed after 6 h. Treatment of rhS100A11 promoted macrophage recruitment in the collagen matrices (n=6 wells, colors represent 2 biological replicates).

### Neutralization of cancer-derived S100A11 in multiple ER+ breast cancer models limits macrophage recruitment

To test whether S100A11 secreted from cancer cells promotes macrophage recruitment, we used both ER+ breast cancer cell lines and a clinically relevant organoid model. First, we characterized secreted S100A11 levels in ER^+^ breast cancer cell lines using ELISA and found that T47D cells secreted a higher concentration of S100A11 than MM330 or BT483 (**Fig. 4A**). Next, we evaluated macrophage recruitment towards cancer cells embedded in a 3D extracellular matrix in this panel of ER+ breast cancer models (**Fig. 4B**). Consistent with the S100A11 secretion findings, we showed that S100A11-high T47D cancer cells recruited a higher number of primary human PBMC-derived macrophages compared to S100A11-low MM330 or BT483 cancer cells (**Fig. 4B**). Using our 3D cancer-macrophage assay, we also visualized the kinetics of macrophages as they infiltrated towards T47D breast cancer cells over a period of 24 h (**Fig. 4C**). To functionally perturb S100A11 during cancer-macrophage coculture, we neutralized S100A11 using a blocking antibody. S100A11 neutralization significantly decreased the number of recruited macrophages in both primary PBMC-derived macrophages and the THP1 macrophage cell line (**Fig. 4D and E**). To evaluate the direct effects of S100A11 neutralization on T47D cancer cells, we compared cancer cell viability after treatment with control (IgG) or anti-S100A11 and found no significant differences (**Supplementary Figure 3**). In addition, we investigated the effects of S100A11 neutralization in a clinically relevant ER breast cancer organoid model (**Fig. 4E**). Macrophage infiltration was reduced in 3D organoids treated with the S100A11 blocking antibody compared to that in the IgG control (45% reduction, p<0.05, **Fig. 4F**). Collectively, our results confirm the bioinformatics findings in primary ER+ breast tumors and highlight the role of S100A11 as a regulator of macrophage recruitment in a 3D microenvironment.

**Figure 4.**
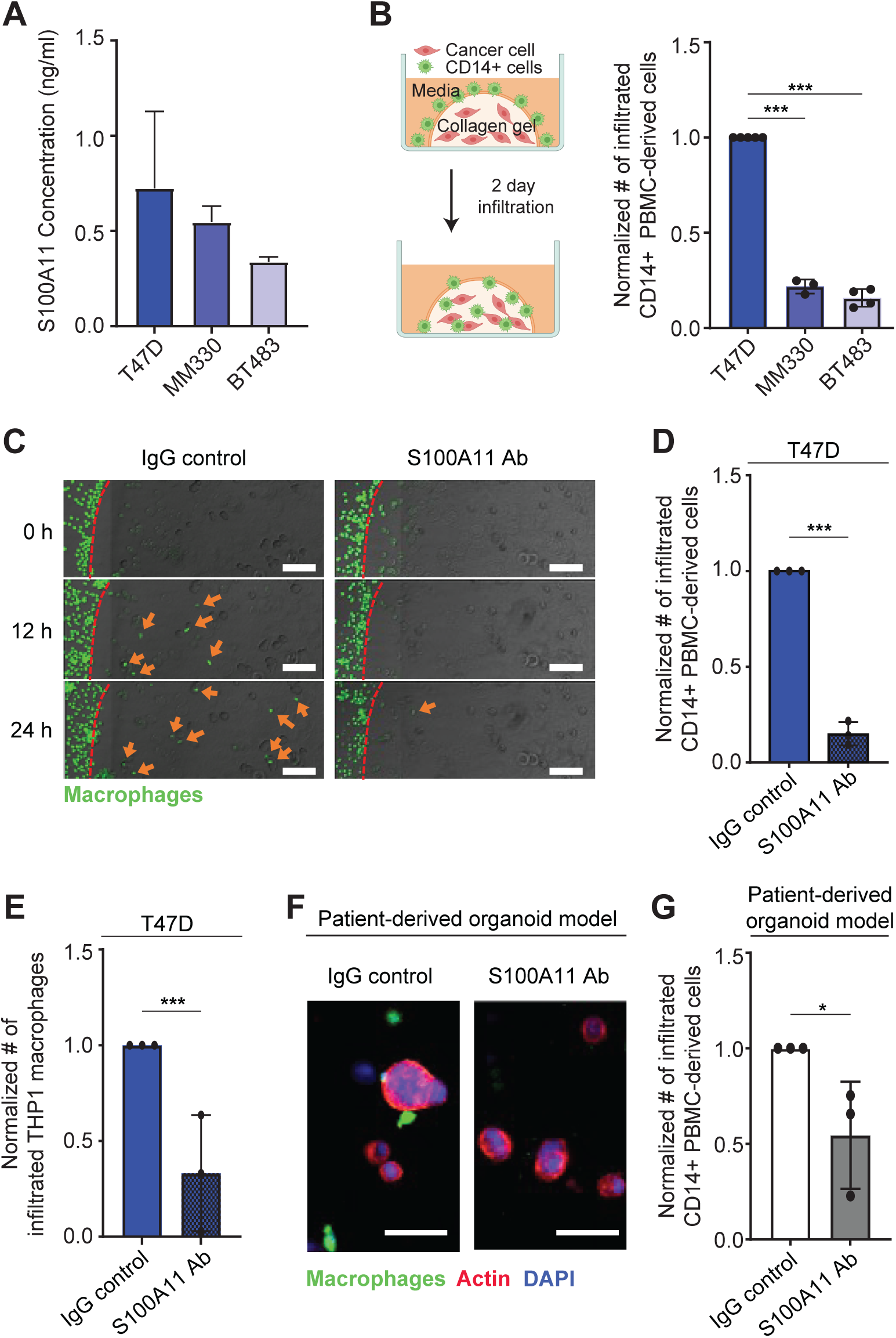
ER+ breast cancer cell secreted S100A11 promotes macrophage recruitment in the 3D collagen matrix. **(A)** Elisa assay to quantify S100A11 secretion level by ER+ breast cancer cell lines showed the highest S100A11 secretion from T47D compared to MM330 or BT483 (n=2 biological replicates). **(B)** 3D collagen droplet experiment to quantify CD14+ PBMC-derived macrophage recruitment to cancer cells. T47D cancer cells recruited highest number of macrophages compared to MM330 or BT483 (biological replicates: n=4 for T47D, BT483; n=3 for MM330). **(C)** Time-lapse images of macrophage infiltration into 3D collagen matrices embedded with T47D for 24 h. Red dotted lines indicate the interface between media and collagen matrices. Orange arrows indicate the infiltrated CD14+ PBMC-derived macrophage. Neutralization of S100A11 by the treatment of S100A11 antibody decreased the recruitment of macrophages. Scale bars, 100 µm. **(D-E)** Normalized number of infiltrated CD14+ PBMC-derived macrophages and THP1 macrophages into the collagen matrices embedded with T47D under the treatment of S100A11 Ab (n=3 biological replicates). **(F)** Visualization of breast cancer organoids embedded in the collagen gel after 24 h in IgG control or S100A11 Ab treatment media. Scale bars, 50 µm. Red: actin staining; Green: CMFDA macrophage staining; Blue: nuclear staining. **(G)** Treatment of S100A11 antibody decreased macrophage recruitment toward breast cancer organoids in collagen matrices (n=3 biological replicates).

## Discussion

A better understanding of cancer-macrophage crosstalk mechanisms is necessary to develop prognostic biomarkers and new therapies to target the pro-tumorigenic tumor ecosystem. Here, we utilized public scRNA-seq datasets to define the ER+ breast cancer TME and identified molecular predictors of macrophage-rich ER+ breast tumors, with a focus on cancer cell-derived factors. S100A11 was the top-ranked gene encoding a secreted factor and was associated with poor patient outcomes. Using a 3D tumor-macrophage experimental platform, we demonstrated that both exogenous and cancer cell-derived S100A11 promoted macrophage trafficking. Neutralization of S100A11 using a blocking antibody reduces macrophage infiltration across multiple macrophages and ER+ breast tumor models. These findings demonstrate the critical role of S100A11 in establishing a pro-tumorigenic macrophage-rich ER+ breast tumor microenvironment and highlight its potential as a therapeutic target.

S100A11 belongs to the S100 family of calcium-binding proteins and is known to regulate cell migration and proliferation via cytoskeletal remodeling in multiple cancer types [23–25]. Consistent with our findings in ER+ breast cancer cells, previous studies have shown that S100A11 is heterogeneously expressed in solid tumors [31, 32]. Overexpression of S100A11 in cervical cancer cells increases their proliferative and migratory abilities [23]. Furthermore, S100A11 has been previously regulates heterotypic cancer-fibroblast crosstalk in the tumor microenvironment [33]. Fibroblast proliferation was increased in co-culture with S100A11-high pancreatic cancer cells, and these fibroblasts promoted tumor growth through a mechanism that was dependent on the S100A11 receptor RAGE [33]. Collectively, these results demonstrate the direct effects of S100A11 on promoting cancer cell pro-tumorigenic functions, as well as its indirect effects via reprogramming stromal fibroblasts in the tumor microenvironment.

S100A11 has not been previously investigated in the context of cancer-macrophage interactions in breast cancer, unlike other S100 family members, such as S100A8 and S100A9, which have been shown to regulate tumor-immune crosstalk [34–38]. Previous transcriptomic and proteomic analyses have shown that S100A11 is overexpressed in breast cancer tissues compared with normal breast tissues [39, 40]. Importantly, the association between high S100A11 gene expression and poor survival outcomes has been previously shown across all breast cancer subtypes [41, 42]. In agreement with our findings on cancer-macrophage crosstalk in ER+ breast cancer, a previous study employed bulk transcriptomics in glioblastoma and found that high expression of S100A11 predicted high infiltration of multiple immune cell types, including macrophages [43]. Another glioblastoma study used scRNA-seq analysis to independently confirm that S100A11-high cancer cells are associated with higher immune cell numbers [44]. Although no previous study has investigated the interaction between cancer cell-derived S100A11 and macrophages, it has been shown that knockdown of S100A11 in cholangiocytes (hepatic epithelial cells) reduced pro-inflammatory reprogramming of macrophages [45]. Taken together, previous investigations of S100A11 in other tumor types support our findings of S100A11-mediated macrophage infiltration in ER breast tumors.

Macrophages represent the most abundant immune cell type in the ER+ breast tumor microenvironment, and there is strong interest in targeting their interactions with cancer cells [21]. Most investigations on cancer-macrophage paracrine factors have focused on well-known chemokines, including CCL2 and CCL5 [13]. Extracellular levels of these chemokines were upregulated in breast tumors compared to normal breast tissue, and treatment with anti-CCL2 and anti-CCL5 blocking antibodies reduced breast cancer dissemination in a zebrafish model [13]. Our findings on cancer cell-driven macrophage recruitment in a 3D matrix are supported by a previous study that employed a 2D Transwell filter to monitor infiltration of the THP1 macrophage cell line towards the ER+ breast cancer cells T47D [46]. As an alternative approach, this study did not evaluate the blockade of cancer-derived ligands but instead showed the potential of IL-1RΑ receptor neutralization [46]. Finally, treatment with an anti-S100A11 antibody suppresses cancer growth in a subset of mesothelioma xenografts [32]. However, no studies have investigated the effects of S100A11 neutralization in breast cancer models.

Nonetheless, our study has some limitations. First, the relatively small sample size imposes constraints on the flexibility of the multivariable modeling. To enable a joint analysis of public datasets, we chose a data integration approach (for detailed information, please refer to the Methods section). It is important to note that each cohort entailed a batch effect due to slight variations in sample processing and differences in scRNA-seq library preparation and sequencing. However, the integration of several datasets has the advantage of mitigating biases related to cell preparation and dissociation, such as preferential liberation of specific cell types during tissue dissociation. Furthermore, our study lacked matching clinical data regarding treatment allocation and treatment responses. This limitation was due to the retrospective nature of the analysis. In addition, the complex signals in the breast tumor microenvironment and the heterogeneous spectrum of macrophage activation pose challenges when analyzing protumorigenic M2-like or antitumorigenic M1-like macrophage states using transcriptomic data [47, 48]. Functional assays that evaluate the effects of tumor-associated macrophage states on cancer cell phenotypes (e.g., growth and invasion) are best suited to address macrophage subpopulation heterogeneity. Hence, it is important to perform future studies on dissecting S100A11-mediated mechanisms of macrophage reprogramming that impact cancer cell phenotypes as well as profiling cell surface marker expression (e.g., pro-tumor: CD163, CD169, CD204, and CD206; anti-tumor: CD80, CD86, iNOS, and HLA-DR) in these macrophages.

In summary, our systems biology approach exploits the heterogeneity of macrophage infiltration patterns in patient tumors and employs 3D tumor models to study the regulatory mechanisms of macrophage recruitment by cancer cells. The identification of therapeutic targets that are enriched in macrophage-rich breast tumors compared with normal breast tissue is critical for developing effective macrophage-directed therapies.

## Materials and Methods

### Publicly available dataset collection

We curated two scRNA-seq human breast cancer datasets encompassing a total of 25 ER+ breast tumor samples. Processed scRNA-seq data from Wu et al. [15] were obtained from the NCBI Gene Expression Omnibus (GEO) under accession number GSE176078, and raw count matrices by Bassez et al. [16] were retrieved from https://lambrechtslab.sites.vib.be/en/single-cell. Gene expression and survival data for ER+ breast cancer patients (n=1,498) were obtained from The Molecular Taxonomy of Breast Cancer International Consortium (METABRIC) [29] through the cBioPortal for Cancer Genomics (http://cbioportal.org) (http://cbioportal.org) [49]. The sample and data are summarized in **Supplementary Table 1**.

### Analysis of scRNA-seq data

#### Data preprocessing and quality control

For the Wu et al. dataset, the matrix.mtx, features.gsv, and barcodes.tsv files were imported into a Seurat object using the Read10X function in the Seurat R package (version 4.3.0) [50]. The count matrix for the Bassez et al. dataset was obtained from the RData file. Subsequently, we filtered the cells exclusively from the ER+ pretreatment patients for further analyses. Cells included in the analysis met the following criteria: they expressed fewer than 6,000 genes, with a minimum of 200 expressed genes and 400 UMI counts. Moreover, these cells exhibited less than 15% reads mapped to mitochondrial gene expression.

#### Batch correction and harmony integration

We applied SCTransform to each Seurat object for data normalization and transformation [51]. SCTransform is a technique designed to mitigate technical variations and alleviate batch effects in scRNA-seq datasets. We merged all the Seurat objects into a single combined dataset to increase the sample size and enhance the statistical power of our analysis. Then, the SCTransform was applied again, regressing the mitochondrial read percentage per cell. Principal component analysis (PCA) was performed on the filtered feature-by-barcode matrix. Uniform Manifold Approximation and Projection (UMAP) embeddings [52] were based on the first 30 principal components. Subsequently, data integration was performed using R package Harmony ver. 1.0 [53] using the function *RunHarmony* to remove batch effects among the samples. The integrated data contained 28,622 genes across 72,079 cell lines.

#### Major cell type identification

The number of dimensions for clustering was chosen based on harmony-embedding clustering. The myeloid (*TYROBP*, *LYZ, CD68*, and *IL1B*) subset was first identified to identify macrophages, monocytes, dendritic cells, and monocytes. The NK/T cell subset was first identified to identify CD4^+^ / CD8^+^ T cells and NK cells. Cells were assigned into one of the following 14 cell types based on the expression of marker genes in each cluster: (i) immune cells (*PTPRC*): B cells (*CD79A*, *MZB1*, *MS4A1*), CD4^+^ T cells (*CD3D*, *CD4, FOXP3*), CD8^+^ T cells (*CD3D*, *CD8A, CD8B*), dendritic cells (*CD40*, *CD1C*, and *ITGAX*), monocyte (*FCGR3Aand CD68*), macrophage (*CD14*, *CD68*, and *FCGR1A*), myeloid (*TYROBP, LYZ,* and *CD68*), mast cells (*CPA3, TPSAB1,* and *TPSB2*) and natural killer (T) cells (*CD3D*^-/+^, *KLRB1, AREG,* and *IL2RB*); (ii) tumor cells (*EPCAM*) or epithelial cancer cells (*KRT18,* and *KRT19*); (iii) Stromal cells (*COL1A1*): myofibroblasts (*ACTA*2), and fibroblasts (*COL1A1 and PDGFRB*); (iv) endothelial cells (*PECAM1*).

#### Macrophage proportion calculation

Following cell type identification, we determined the macrophage proportion for each patient and arranged them in ascending order based on this fraction. Similarly, we computed the fraction of each cell type in individual patients and utilized these proportions to assess the correlation between different cell types. Spearman correlation was calculated, and the correlation coefficient (rho) was visualized in a heatmap. We calculated the mean gene expression in individual patients and calculated the Spearman correlation between gene expression and macrophage proportions in different cell types.

### Survival analysis

METABRIC ER+ RNA-seq and clinical data were used for survival analysis. Patients with the top 25% and bottom 25% S100A11, LY6G6C, and PDIA4 expression were defined as high and low groups, respectively. A Cox proportional hazards regression model was fitted against the groups, and Kaplan–Meier survival curves were drawn using the R package survival (ver. 3.5-5) (https://cran.r-project.org/web/packages/survival/)

### Human protein atlas

The Human Protein Atlas (https://www.proteinatlas.org) is a public resource that extracts information, including images of immunohistochemistry (IHC), protein profiling, and pathologic information, from specimens and clinical material from cancer patients to determine global protein expression [30]. Here, we compared the protein expression of S100A11 in tumor and normal breast tissues using IHC.

### Cell culture

The T47D and BT483 cell lines were cultured in RPMI medium, and the MM330 cell line was cultured in DMEM/L15 media supplemented with 10% FBS and 1% penicillin-streptomycin. Human peripheral blood mononuclear cells (PBMCs) were isolated from healthy donors and CD14+ cells were isolated using CD14 microbeads (Cat# 130-050-201, Miltenyi Biotec). THP1 cells were cultured in RPMI medium supplemented with 10% FBS and 1% penicillin-streptomycin. THP1 cells were treated with 10 ng/ml phorbol 12-myristate 13-acetate (PMA) for 24 h to differentiate into macrophages. Patient-derived breast cancer organoids (BO#129, ER+/PR+ breast tumor) were generated at the Institute of Precision Medicine (Pitt/UPMC) and cultured in 40 µL of Cultrex Basement Membrane Extract (Cat. 343200101, R&D Systems™) in 24 well plate in organoid medium (recipe and culture procedures in) [54].

### Live imaging and migration assay

CD14+ monocytes were seeded in 96 well plate (50,000 cells per well) and treated with 25 ng/ml macrophage colony-stimulating factor in RPMI supplemented with 10% FBS and 1% penicillin-streptomycin for six days to differentiate them into macrophages. For cell migration analysis, macrophages were stained with CellTracker Green 5-chloromethylfluorescein diacetate (CMFDA) dye (Cat. C7025, Invitrogen™) and imaged every 2 min for 60 min using a Zeiss LSM700 confocal microscope. Migration trajectories and average speeds were quantified using the Trackmate plugin in ImageJ.

### 3D Inverted invasion assay in the presence of S100A11 concentration gradients

Following CD14+ differentiation into macrophages in 96 well plates, a 2 mg/ml collagen matrix was formed on top of the cells by polymerization of collagen type I solution for 45 min at 37°C. Next, 100 µL of medium with different concentrations of recombinant S100A11 was added to establish a concentration gradient. We imaged five z-stacks (20 µm interval) every 1 hour to monitor macrophage recruitment in the 3D collagen matrix. The number of macrophages in each z-section was quantified using Trackmate in ImageJ software.

### 3D tumor-macrophage droplet infiltration experiment

ER+ breast cancer cell lines or patient-derived cancer organoids were mixed with 2 mg/ml collagen type I solution (1E6 cells/ml concentration) and seeded in each well of 96 well plate. After 30 min, the solution was polymerized into a collagen matrix and 50k of CMFDA-stained PBMC-derived CD14+ cells were seeded in 100 µL of media per well. After 48 h of tumor-macrophage coculture, the plate was imaged using a confocal microscope to assess the number of macrophages recruited inside the collagen matrices.

### Statistical analysis and visualization

Statistical analyses and visualizations were performed using R. The statistical methods used for each analysis are described in the text and the figure legends. Statistical significance was set *p*-value < 0.05. Graphs were generated using R package ggplot2 (ver. 3.3.6) and ggpubr (ver. 0.4.0), Ggrepel, gridExtra (ver. 2.3), ComplexHeatmap (ver. 2.16.0) [55], and Fgsea (ver. 1.26.0). Violin plots of single-cell data were drawn using VlnPlot in the Seurat R package (ver. 4.3.0) [50]. We used ggplot2 and geom_violin to draw a violin plot for compact display of a continuous distribution of pathway activities in single cells. Geom _violin is a blend of geom_boxplot and geom_density, which computes kernel density estimates. All boxplots report the 25% (lower hinge), 50%, and 75% quantiles (upper hinge). The lower (upper) whiskers indicate the smallest (largest) observation greater (less) than or equal to the lower (upper) hinge −1.5^ interquartile range (IQR) (+1.5^IQR) as default in the *geom_boxplot* function ggplot2.

### Data availability

The code used to analyze the integrated scRNA-seq in this study is available in the open access database at https://codeocean.com/capsule/6931150/tree. The raw data underlying Figs 4-5 are presented in the Supplementary Data.

## Abbreviations

APOD: apolipoprotein D
CCL: chemokine ligand family
CMFDA: chloromethylfluorescein diacetate
DCs: dendritic cells
ECCs: epithelial cancer cells
ER+: estrogen receptor-positive
GEO: Gene Expression Omnibus
HPA: Human Protein Atlas
HR: hazard ratio
IHC: immunohistochemically
IQR: interquartile range
METABRIC: Molecular Taxonomy of Breast Cancer International Consortium
NK: natural killer
PBMC: peripheral blood monocyte
PCA: principal component analysis
PMA: phorbol 12-myristate 13-acetate
scRNA-seq: Single-cell RNA-seq
TME: tumor microenvironment
Tregs: regulatory T cells
UMAP: uniform manifold approximation and projection

## Acknowledgments

We would like to thank the authors of [15, 16] for sharing their data on breast cancer scRNA-seq data. The data analyses in this research were supported by the University of Pittsburgh Center for Research Computing (CRC) and the Extreme Science and Engineering Discovery Environment (XSEDE), which is supported by the National Science Foundation grant number OCI-1053575. Specifically, it used the Bridges2 system, which is supported by NSF award number ACI-1445606 at the Pittsburgh Supercomputing Center (PSC).

## Contributions

HUO and IKZ designed and supervised the study, analysis, and writing of the manuscript. SL performed the bioinformatics analysis and helped write the manuscript. YC performed experimental validation and helped write the manuscript. YL and RL performed experiments. DB and PM generated the organoid models. AL and SO contributed to discussions on project design, provided resources, and provided helpful suggestions for revisions of the manuscript. All authors have read and agreed to the publication of this manuscript.

## Ethics declarations

### Ethics approval and consent to participate

Not applicable.

### Consent for publication

Not applicable.

